# Coherent representations of subjective spatial position in primary visual cortex and hippocampus

**DOI:** 10.1101/235648

**Authors:** Aman B. Saleem, E. Mika Diamanti, Julien Fournier, Kenneth D. Harris, Matteo Carandini

**Affiliations:** UCL Institute of Ophthalmology, University College London, London, EC1V9EL, UK; Department of Experimental Psychology, University College London, London, WC1H 0AP, UK; CoMPLEX, Department of Computer Science, University College London, London, WC1E 6BT, UK; UCL Institute of Neurology, University College London, London, WC1N 3BG, UK

**Author notes:** Equal contribution (Aman B. Saleem, E. Mika Diamanti). These authors jointly supervised this work.

## Abstract

A major role of vision is to guide navigation, and navigation is strongly driven by vision^1-4^. Indeed, the brain’s visual and navigational systems are known to interact^5, 6^, and signals related to position in the environment have been suggested to appear as early as in visual cortex^6, 7^. To establish the nature of these signals we recorded in primary visual cortex (V1) and in the CA1 region of the hippocampus while mice traversed a corridor in virtual reality. The corridor contained identical visual landmarks in two positions, so that a purely visual neuron would respond similarly in those positions. Most V1 neurons, however, responded solely or more strongly to the landmarks in one position. This modulation of visual responses by spatial location was not explained by factors such as running speed. To assess whether the modulation is related to navigational signals and to the animal’s subjective estimate of position, we trained the mice to lick for a water reward upon reaching a reward zone in the corridor. Neuronal populations in both CA1 and V1 encoded the animal’s position along the corridor, and the errors in their representations were correlated. Moreover, both representations reflected the animal’s subjective estimate of position, inferred from the animal’s licks, better than its actual position. Indeed, when animals licked in a given location – whether correct or incorrect – neural populations in both V1 and CA1 placed the animal in the reward zone. We conclude that visual responses in V1 are tightly controlled by navigational signals, which are coherent with those encoded in hippocampus, and reflect the animal’s subjective position in the environment. The presence of such navigational signals as early as in a primary sensory area suggests that these signals permeate sensory processing in the cortex.

---

To characterise the influence of spatial position on the visual responses of area V1 we took transgenic mice expressing the calcium indicator GCaMP6 in excitatory cells and placed them in a 100 cm corridor in virtual reality (VR; Figure 1**a**). The corridor had four prominent landmarks, spaced 20 cm apart: a grating and a plaid, and then again a grating and a plaid. The repetition of landmarks created two visually-matching segments of the corridor, separated by 40 cm (Figure 1**a, b**; Supplementary Figure 1). We identified V1 based on the retinotopic map of the cortical surface, measured using wide-field imaging^8^ (Figure 1**c**). We then pointed a two-photon microscope on regions in medial V1, focusing our analysis on neurons with receptive field centres >40° azimuth (Figure 1**c**), which were driven as the mouse passed the landmarks. As expected, given the repetition of visual scenes in the two segments of the corridor, some V1 neurons had a response profile with two equal peaks spaced 40 cm apart (Figure 1**d**). Other V1 neurons, however, responded very differently to the same visual stimuli in the two segments (Figure 1**d**). These results indicate that visual activity in V1 can be strongly modulated by the animal’s spatial position in an environment.

**Figure 1:**
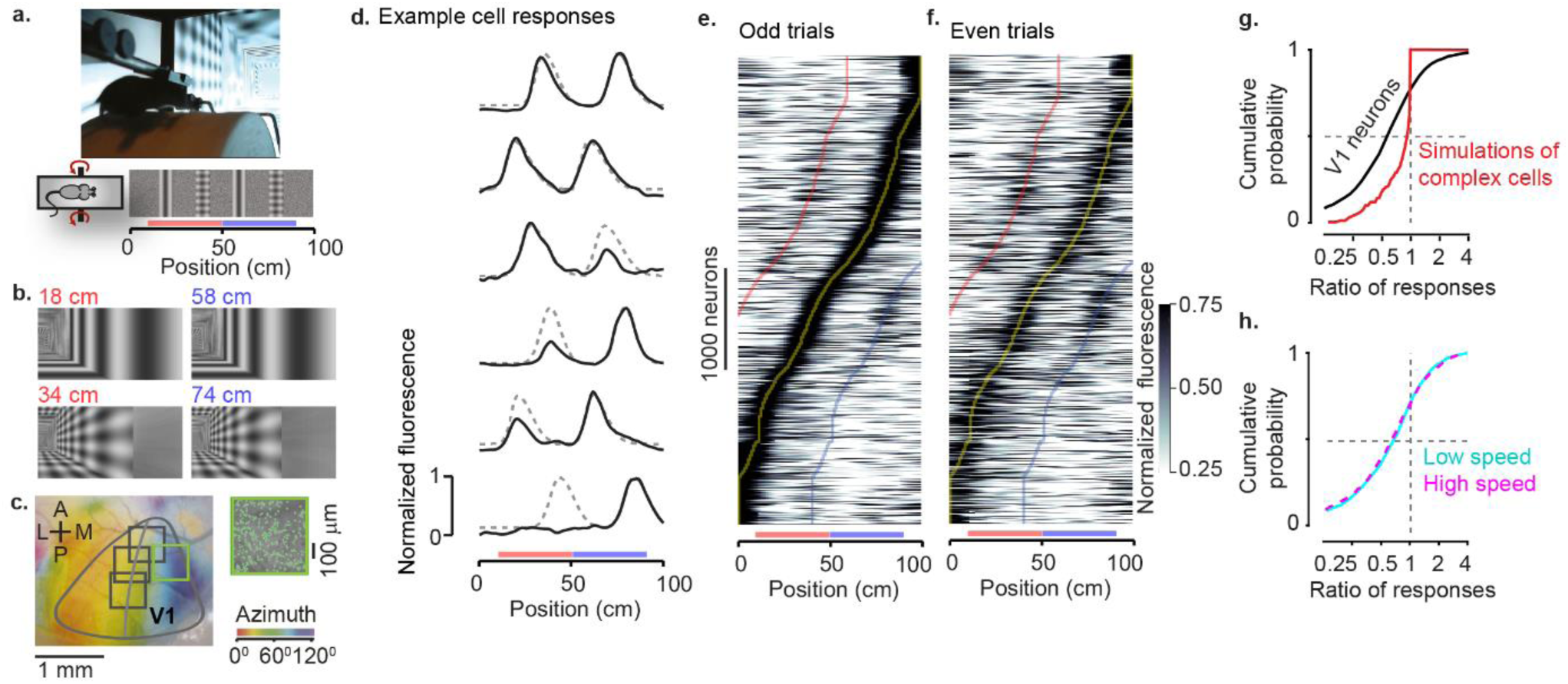
Responses in the primary visual cortex (V1) are modulated by spatial context. **(a)** Mice ran on a cylindrical treadmill to navigate a virtual corridor presented on three visual displays. The corridor had two landmarks (a grating and a plaid) that repeated after 40 cm, creating two visually-matching segments (*red* and *blue bars*). **(b)** Example screenshots, showing the visual similarity of the virtual corridor at two pairs of positions spaced 40 cm apart. **(c)** Example retinotopic map of the cortical surface acquired with widefield calcium imaging. The border of V1 is defined by inversion of the retinotopic map. *Squares* denote the field of view in two-photon imaging sessions targeted to medial V1 in an animal (field of view with *green frame* is shown in the *inset*). Within these sessions we analysed responses from neurons with receptive field centre > 40^°^ azimuth *(curve).* **(d)** Normalized response as a function of position in the virtual environment for six example V1 neurons. *Dotted lines* show predictions if the responses were identical in the two segments of the corridor (*red* and *blue bars*). **(e)** Normalized response as a function of distance in the virtual environment obtained from odd trials only, for V1 neurons with receptive field centres >40^°^ azimuth and activity significantly modulated by position in the corridor (4,958 of 8,610 neurons). Neurons are ordered based on the position of their maximum response. *Yellow, red* and *blue lines* indicate position of maximum +/-40 cm. *Red* and *blue bars* are as in **a**. **(f)** Same as **e**, for the half of the data (even trials) that were not used to order the responses. Sequence and scaling of the responses are the same as in **e**. **(g)** Cumulative distribution for the ratio of secondary response (40 cm away from peak response, *red* or *blue line* in **f)** divided by peak response (*yellow line* in **f)**, derived from even trials *(black).* For comparison, the *red* curve shows the same ratio of responses obtained from simulations of Complex cells with purely visual responses. **(h)** Same as in **g** after stratifying the data by running speed. The two curves corresponding to low *(cyan)* and high *(purple)* speeds overlap and appear as a single dashed curve.

This modulation of visual responses by spatial position occurred in the majority of V1 neurons (Figure 1**e-g**). We imaged the activity of 8,610 V1 neurons across 18 sessions in 4 mice. We selected neurons (n = 4,958) with receptive field centres >40^°^ azimuth and activity significantly modulated by position in the corridor, and divided the trials in half, using the odd trials to find the position where each neuron fired maximally. The resulting representation reveals a striking preference of V1 neurons for spatial position, with most neurons giving much stronger responses in one position than in the visually matching position 40 cm away (Figure 1**e**). To quantify this preference while avoiding any circularity, we used the data obtained in the other half of the trials (even trials). These data showed that the preference for spatial position was robust (Figure 1 **f**). Among the neurons that responded when the animal traversed the two visually-matching segments (n = 2,422), the responses to the landmarks 40 cm from the preferred position were substantially smaller than the responses at the preferred position (Figure 1**g**; Supplementary Figure 2): the median ratio of their responses was 0.61 ± 0.31 (± m.a.d.). Modulation of visual responses by spatial position therefore reflected a reliable and widespread preference of individual V1 cells for one of the two locations.

The modulation of V1 responses by spatial position could not be explained by classical visual responses, or by deviations in pupil position and diameter, running speed or reward. As expected, applying a model of purely visual responses on the sequences of images (using a simulation of V1 complex cells, Supplementary Figure 3) generated ratios of responses that were very close to 1 (0.97 ± 0.17). The preference for spatial position could not be explained by deviations in pupil position or size as the ratio of responses was strongly biased towards low values even in sessions with steady eye position or pupil size (Supplementary Figure 4). Moreover, the modulation of V1 responses by spatial position could not be explained by variations in the animal’s speed. Given that V1 neurons are influenced in diverse ways by running speed and visual speed^9-12^, they might respond differently in the two segments of the corridor based on speed differences. To control for this potential effect, we stratified the data in three groups according to instantaneous running speed (low, medium, or high; Supplementary Figure 5), and estimated each neuron’s preferred position based on medium speed. We then calculated the ratio of responses for low and high speeds, and found them to be identical (Figure 1**h**). The spatial modulation of visual responses was also independent of the presence of a reward: in some sessions mice ran freely in the absence of a reward, and the ratio of responses remained significantly different from the simulations of purely visual responses (Supplementary Figure 6).

Having established that V1 responses are modulated by spatial position, we next asked whether the underlying signals reflect the spatial position encoded in the brain’s navigational systems and the animal’s subjective estimate of position (Figure 2). We recorded neuronal activity simultaneously in V1 and hippocampal area CA1 using two 32-channel extracellular electrodes in four wild-type mice (Figure 2**a**). To gauge their subjective estimate of position, we trained the mice to lick a water spout upon reaching a specific region of the VR corridor to receive a water reward (Figure 2**b**; Supplementary Movie 1, Supplementary Figure 7). All four mice learned to perform this task with over 80% accuracy. To ensure that the task required vision and could not be solved simply by counting steps, we introduced random intervals between consecutive runs through the corridor, and also tested that the animals could still perform the task when we changed the gain relating wheel rotation to progression in the VR corridor (Supplementary Fig 7**f, g**). Moreover, performance was reduced when the contrast was lowered, thus confirming that the animal used vision to perform the task (Supplementary Figure 7**c-e**).

**Figure 2:**
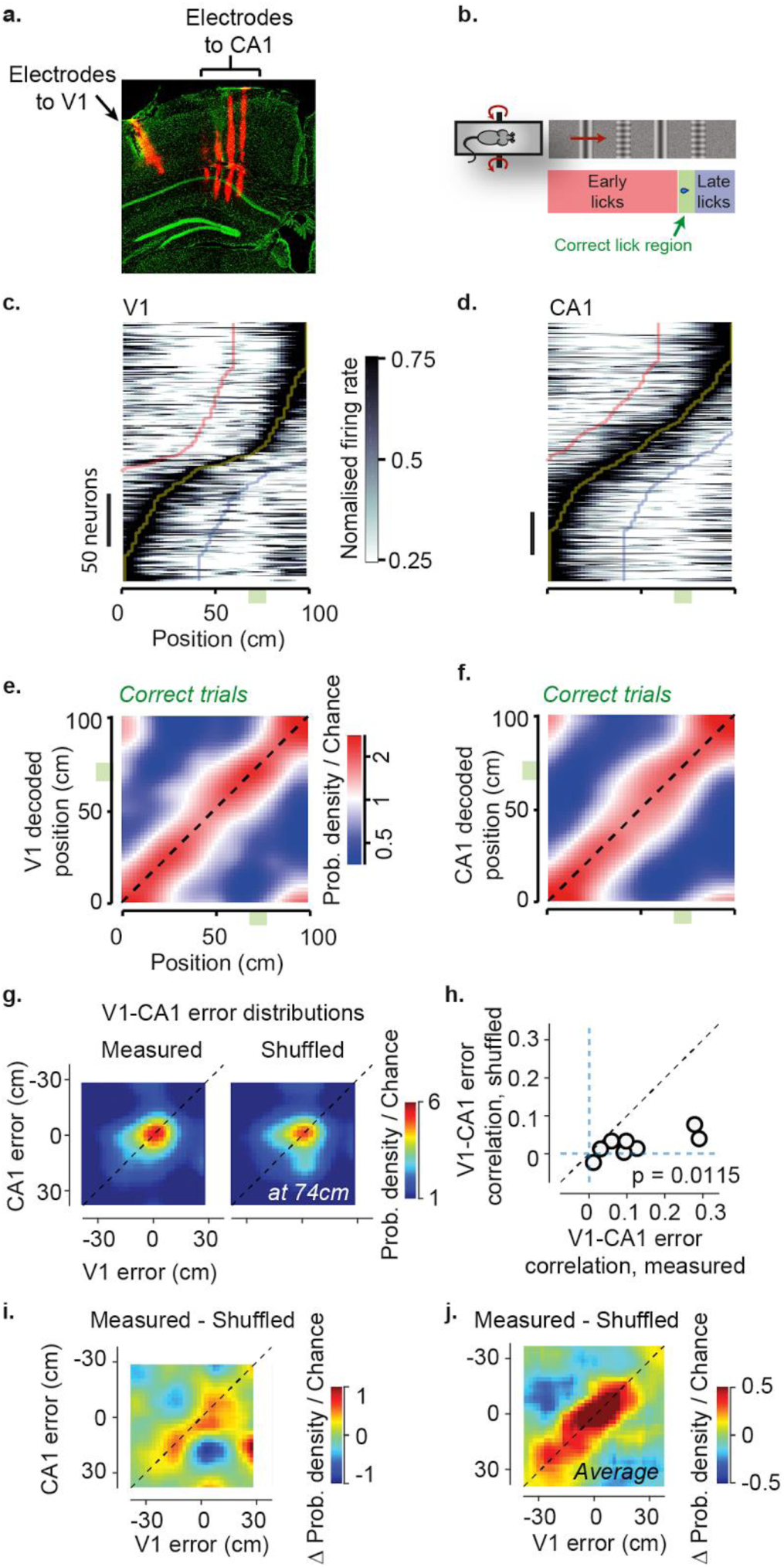
V1 and CA1 neural populations represent spatial positions in the virtual corridor and make correlated errors. **(a)** Example of reconstructed electrode tracks. Multi-electrode arrays were dipped in DiI before insertion *(red); green* shows cells labelled by DAPI. *Panel* shows one array (with four shanks) in the CA1 layer of the Hippocampus together with a second electrode track of one shank aimed at the primary visual cortex (V1). **(b)** In the task, water was delivered when mice licked in a reward zone (indicated by *green shaded area* in bottom of **a**). (**c**) Normalized activity as a function of position in the virtual corridor, for V1 neurons (266 cells, 8 sessions). Cells are ordered based on the position of their maximum response. *Yellow, red* and *blue lines* indicate position of maximum +/- 40 cm. (**d**) Similar plot for CA1 place cells (334 cells, 8 sessions). (**e**) Density map showing the distribution of the position decoded from the firing of simultaneously recorded V1 neurons (y-axis) as a function of the animal’s actual position (x-axis). The distribution of decoded positions was averaged across all recording sessions (n = 8), considering only correct trials. The red diagonal stripe indicates accurate estimation of position from the population. (**f**) Similar plot for CA1 neurons. (**g**) Density map showing the joint distribution of position decoding error from V1 and CA1 at an example position (left), together with a similar analysis on data shuffled preserving the correlation of run speed and position (right). **(h)** Correlation coefficient of decoding errors in V1 and CA1 for each recording session, against similar analysis of shuffled data. Correlations are significantly above shuffling control (p=0.0198, Student t-test). **(i)** Difference between density map for decoding errors from simultaneously recorded V1 and CA1 populations and shuffling control, for the example in **g**. **(j)** Difference between joint density map of V1 and CA1 position, and shuffled control, averaged over all positions and recording sessions (n = 8).

Many recorded neurons in both the visual cortex and hippocampus had place-specific response profiles and faithfully represented the position of the mouse in the environment (Figure 2**c-f**). Consistent with our previous observations with two-photon imaging, V1 neurons displayed a non-visual modulation of their response profiles, i.e. they responded more strongly in one of the two visually-matching segments of the corridor (Figure 2**c**). In turn, hippocampal CA1 neurons exhibited place fields^^4, 13-15^^, responding in a single corridor location (Figure 2**d**). Responses in both V1 and CA1 encoded the position of the mouse in the environment, with no ambiguity between the two visually-matching segments. Indeed, an independent Bayes decoder^16^ was able to read out the animal’s position from the activity of neurons recorded from V1 (33 ± 17 neurons per session, n = 8 sessions; Figure 2e) or from CA1 (42 ± 20 neurons per sessions; n = 8 sessions; Figure 2**f**).

Furthermore, when visual cortex and hippocampus made errors in estimating the mouse’s position, these errors were correlated with each other (Figure 2**g-h**). The distributions of errors in position decoded from V1 and CA1 both peaked at zero (Figure 2**g**), but were significantly correlated (Figure 2**h**; ρ = 0.125, p = 0.0129, Student t-test). Variations in speed affect responses of both V1^9, 10^ and CA1^16-19^, so it is important to test whether this correlation arises from a common modulation of both regions by running speed. We therefore shuffled the data between trials in a manner that preserves the relationship between speed and position. After shuffling, the correlation dramatically decreased: V1 and CA1 decoding errors had a correlation of only 0.022, significantly less than the observed moment-by-moment correlation of 0.125 (p = 0.0115; Figure 2**g, h**). To estimate how V1 and CA1 errors covary regardless of position and speed, we subtracted the joint distributions obtained from the original dataset and from the shuffled one (Figure 2**i, j**). The residual showed a clear distribution of decoding errors along the diagonal, indicating that V1 and CA1 representations are more correlated than expected from common speed modulation. Thus, on a moment-by-moment basis V1 carries an estimate of the animal’s position that is consistent with the estimate in CA1.

We next asked whether the positional signals carried by CA1 and V1 relate to the animal’s subjective estimate of position (Figure 3**a-f**). CA1 activity is influenced by the ongoing performance of navigation tasks^20-23^, and may reflect the animal’s subjective position more than actual position^21, 23, 24^. Having trained the mice to selectively lick the spout in the reward zone, we could assess their subjective estimate of position from the location of their licks. We divided trials into three groups: *Early* trials when too many licks (usually 4-6) were before the reward zone, causing the trial to be aborted; *Correct* trials when one or more licks occurred in the reward zone; and *Late* trials when the mouse missed the reward zone and licked afterwards. To understand how V1 and CA1 neural representations of space related to this behaviour, we trained the Bayesian decoder on the population activity measured in *Correct* trials, and analysed the likelihood of decoding different positions in the three types of trials. Decoding performance in *Early* and *Late* trials showed systematic deviations from the diagonal (where decoded position is veridical). In *Early* trials, V1 and CA1 tended to overestimate the animal’s progress along the corridor (a deviation above the diagonal in Figure 3**a, d**). In *Late* trials, conversely, they tended to underestimate it (a deviation below the diagonal in Figure 3**b, e**). Accordingly, the probability of being in the reward zone peaked before the actual reward zone in *Early* trials and after it in *Late* trials, and this was the case not just for CA1 but also for V1 (Figure 3 **c, f**). These consistent deviations suggest that the representations of position in V1 and CA1 are correlated with the animal’s decisions to lick and thus likely reflect the animal’s subjective estimate of position.

**Figure 3:**
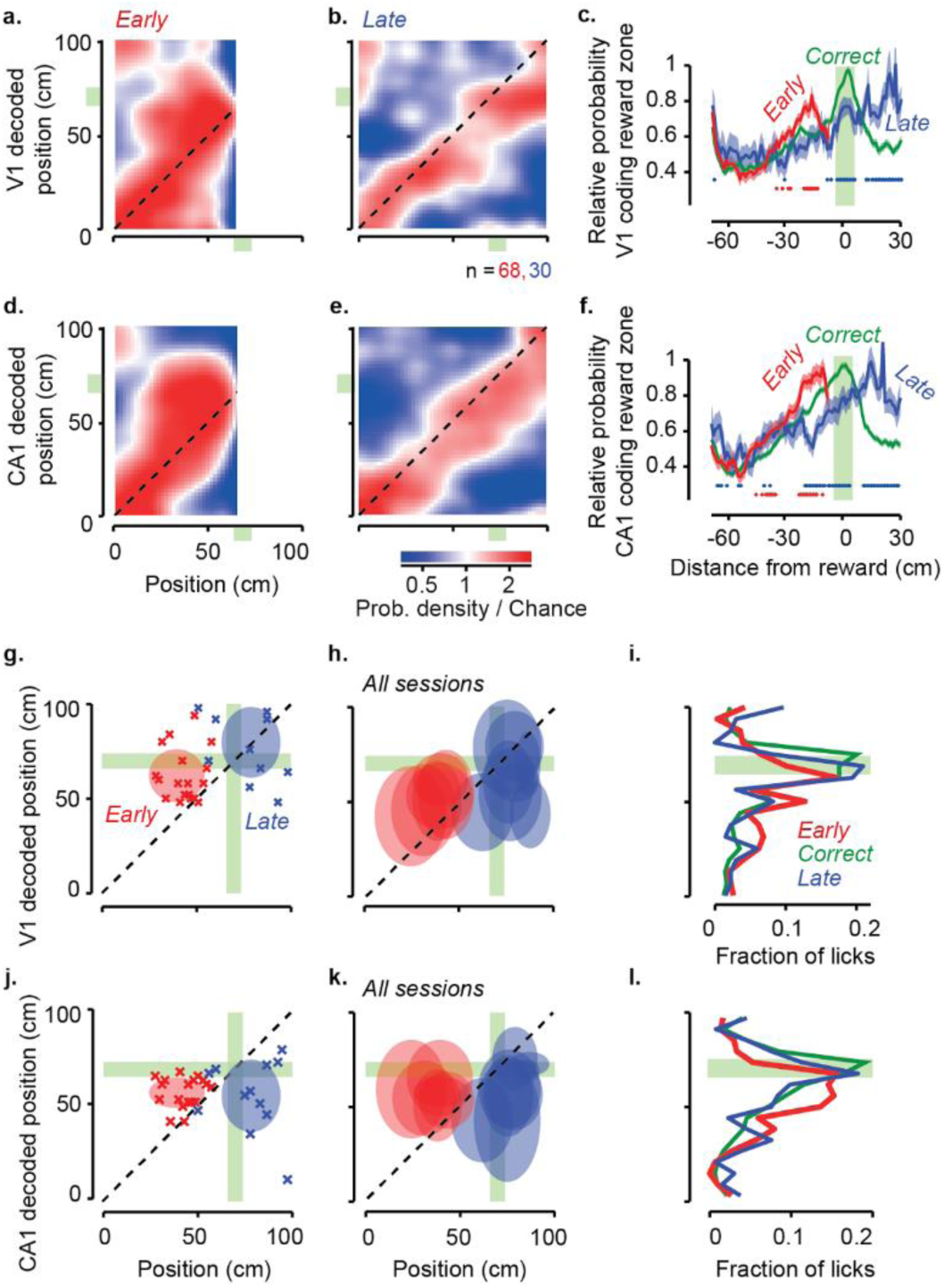
Positions encoded by visual cortex and hippocampus are correlated with animal’s spatial decisions. **(a)** Density map showing the distribution of positions decoded from the V1 population (y-axis), as a function of the animal’s actual position (x-axis) on trials where the mice licked *early*. The decoder was trained on a separate set of trials where mice licked in the correct position. (**b**) Same plot for trials where the animal licked late. (**c**) The average decoded probability for the animal to be in the reward zone, as a function of distance from the reward on *Early (red), Correct (green)* and *Late (blue)* trials. The red curve peaks before the animal actually reaches the reward zone, while the blue curve peaks after, indicating that V1 population activity represents subjective, rather than actual position at the time of licking on early and late trials. The probability was normalized relative to the probability of being in the reward zone in the correct trials. *Red dots*: positions where the decoded probability of being in the reward zone differed significantly between *Early* and *Correct* trials (p<0.05, two-sample t-test). *Blue dots*: significant difference between *Correct* and *Late* trials. (**d-f**) Same as **a-c**, for decoding using the population of CA1 neurons. **(g)** Position decoded from V1 activity as a function of actual position of the animal. Each cross shows the position of when the animal licked for a reward on *Early* trials *(red crosses)* or *Late* trials *(blue crosses)* on an example session. Note that *Late* trials can include some early licks. The distributions of all the error trial licks during the example session (mean ± s.d) are shown as *shaded regions* for *Early (red)* and *Late (blue)* trials. The reward zone is indicated as a *light green shaded region.* **(h)** The distributions of positions decoded from V1 activity as a function of actual position of the animal for all error trial licks across all recording sessions (n = 8). **(i)** Fraction of licks as a function of distance from reward location in decoded positions decoded from V1 activity. **(j-l)** Same as **g-i**, for CA1 neurons.

The timing of the licks provides an opportunity to gauge when the mouse’s subjective estimate of position lies in the reward zone. If activity in V1 and CA1 reflects subjective position, it should place the animal in the reward zone whether the animal correctly licked in that zone or it licked earlier or later. This prediction is borne out by the data (Figure 3**g-l**). In the VR environment, we had a precise measurement of the times when the animal licked. We could therefore decode activity in V1 and CA1 at those times, and compare the decoded position to the animal’s actual position. When plotted as a function of actual position, by definition, the distributions of licks in *Early*, *Correct*, and *Late* trials were distinct (Figure 3**g, h** and **j, k**). However, when plotted as a function of decoded position (with a decoder trained only on *Correct* trials) these distributions came into register over the rewarded zone, whether the decoding was done from V1 activity (Figure 3**g-i**) or from CA1 activity (Figure 3**j-l**). Thus, when animals licked for a reward, the activity of both V1 and CA1 signalled that position was in the reward zone.

Taken together, these results indicate that the visual responses of V1 are modulated by the same spatial signals represented in hippocampus, and that these signals reflect the animal’s subjective position in the environment. These signals may become stronger as animals become increasingly familiar with the environment^6, 7^, perhaps contributing to the changes in V1 responses to visual stimuli seen as animals learn behavioural tasks^25-27^. However, we also observed spatial modulation in animals that freely ran the environment without reward (Figure 1; Supplementary Figure 6), suggesting that even incidental learning of the spatial features of the environment, in the absence of a task, is sufficient to modulate V1 responses.

The network of connections underlying this modulation of visual responses by subjective spatial position is yet unknown. While the primary visual cortex and hippocampus are not directly connected, both feed-forward and feedback connections (possibly through higher visual areas and retrosplenial or rhinal cortices) may convey spatial information between the two regions^28, 29^. Outside the hippocampal formation, spatial signals have been reported in the retrosplenial, parietal and prefrontal cortex^30, 31^. Our data show that spatial signals related to an animal’s own estimate of position appear as early as in a primary sensory cortex. This result indicates that the mouse cortex does not keep a firm distinction between navigational and sensory systems: rather, spatial signals permeate cortical processing.

## Acknowledgements

We thank Bilal Haider and other members of the lab for helpful discussions, and Neil Burgess for comments on the project. We thank Charu Reddy for help with histology, and Sylvia Schroeder and Michael Krumin with two-photon imaging. ABS is a recipient of the Sir Henry Dale Fellowship, awarded by Wellcome Trust / Royal Society (grant 200501). JF was supported by Human Frontier Science program (HFSP) and EC | Horizon 2020 (EU Framework Programme for Research and Innovation). MC and KDH are jointly funded by the Wellcome Trust (grant 095668 and 095669) and by the Simons Collaboration on the Global Brain (grant 325512). MC holds the GlaxoSmithKline/Fight for Sight Chair in Visual Neuroscience.

## Author contributions

All authors contributed to the design of the study. ABS & MD carried out the experiments and analysed the data with JF. ABS, MD, JF, KDH and MC wrote the paper.

## Supplementary Figures

**Supplementary Movie 1: Movie of animal performing the task in virtual reality.**

**Supplementary Figure 1:**
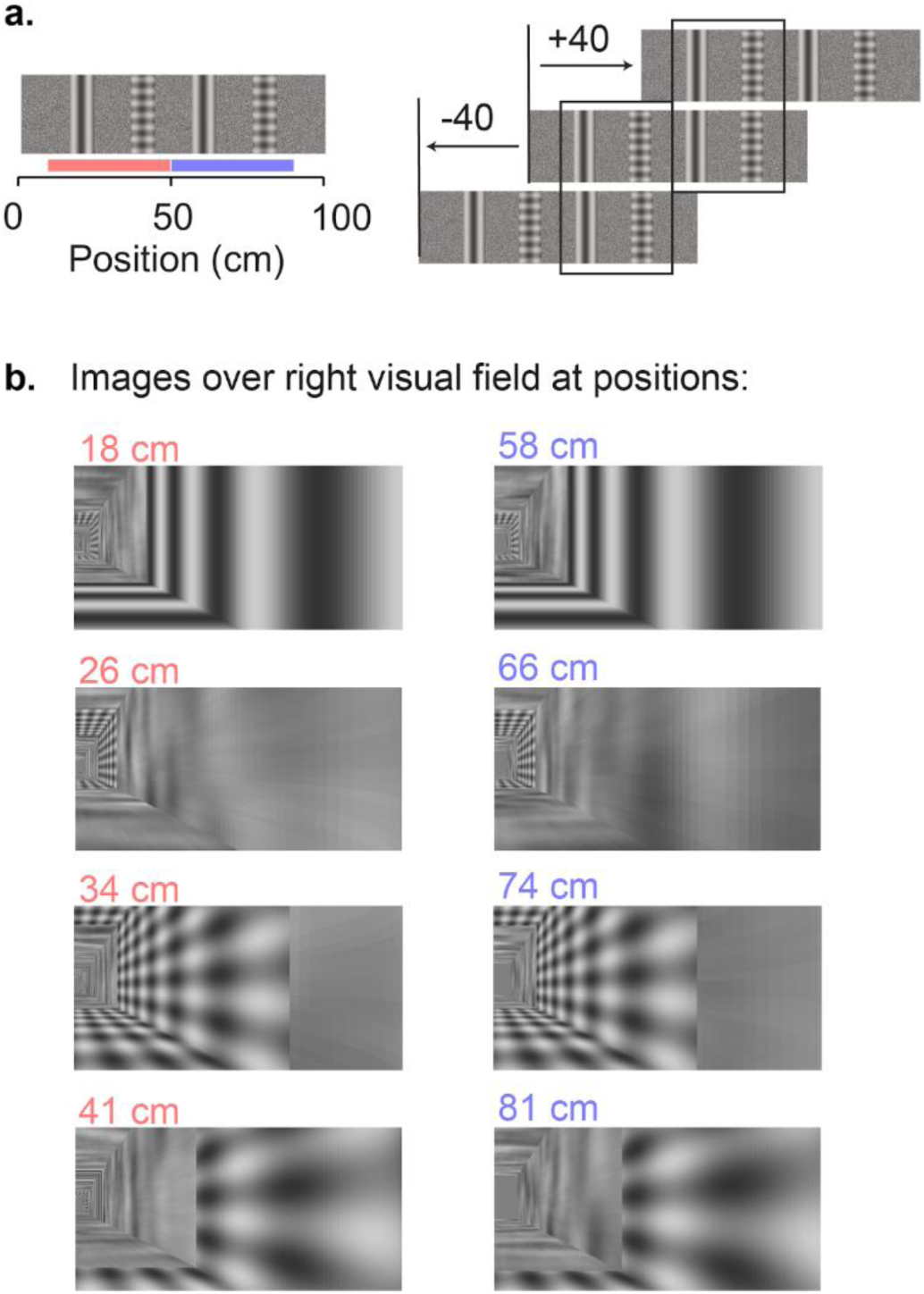
Design of virtual environment with two visually matching segments. **a,** The virtual corridor had four prominent landmarks. Two landmarks (grating and plaid) were repeated at two positions, 40cm apart creating two visually matching segments in the room, from 10 cm to 50 cm and from 50 cm to 90 cm (indicated by *red* and *blue* bars in the *left panel*), as illustrated in the *right panel*. **b,** Example screenshots of the right visual field displayed in the environment when the animal is at different positions. Each row displays screen images at positions 40cm apart.

**Supplementary Figure 2:**
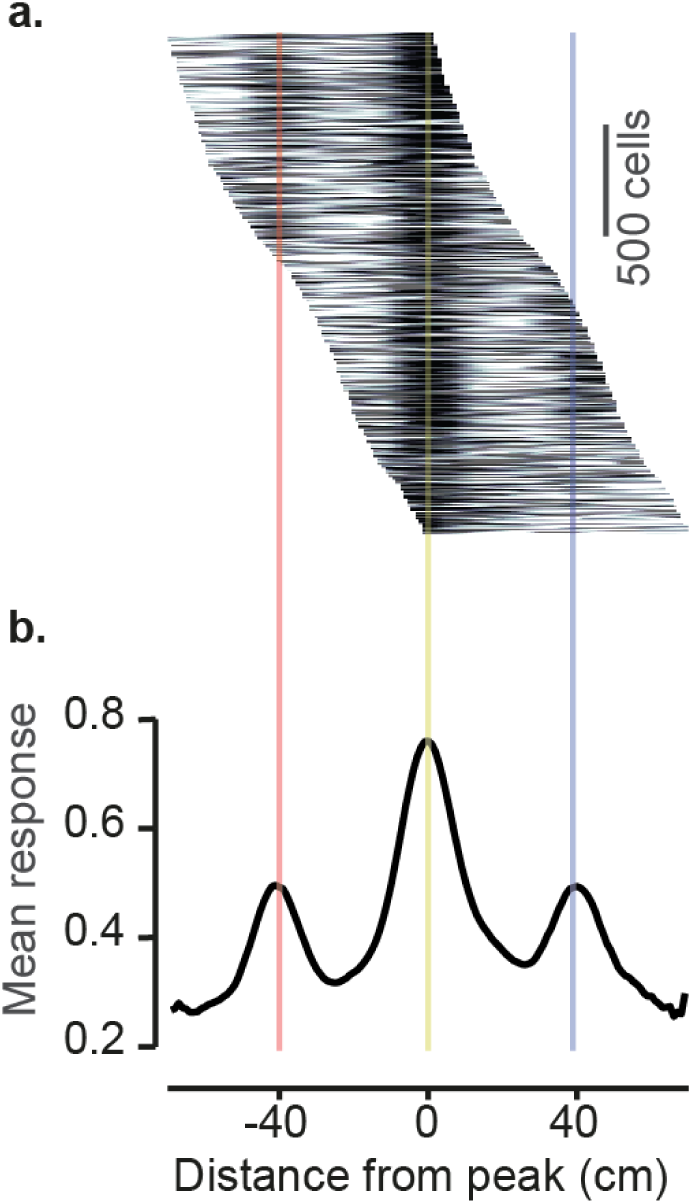
Spatial averaging of visual cortical activity confirms the difference in response between visually matching locations. **a,** Mean response of V1 neurons as a function of the distance to the peak response, as obtained from even trials (2,422 cells with peak response between 15 and 85 cm along the corridor). The position of the peak response was estimated from the other half of the trials (odd trials). **b,** Population average across responses shown in a. Lower values of the side peaks compared to central peak indicates strong preference of V1 neurons for one segment of the corridor over the other visually-matching segment (40 cm away from peak response).

**Supplementary Figure 3:**
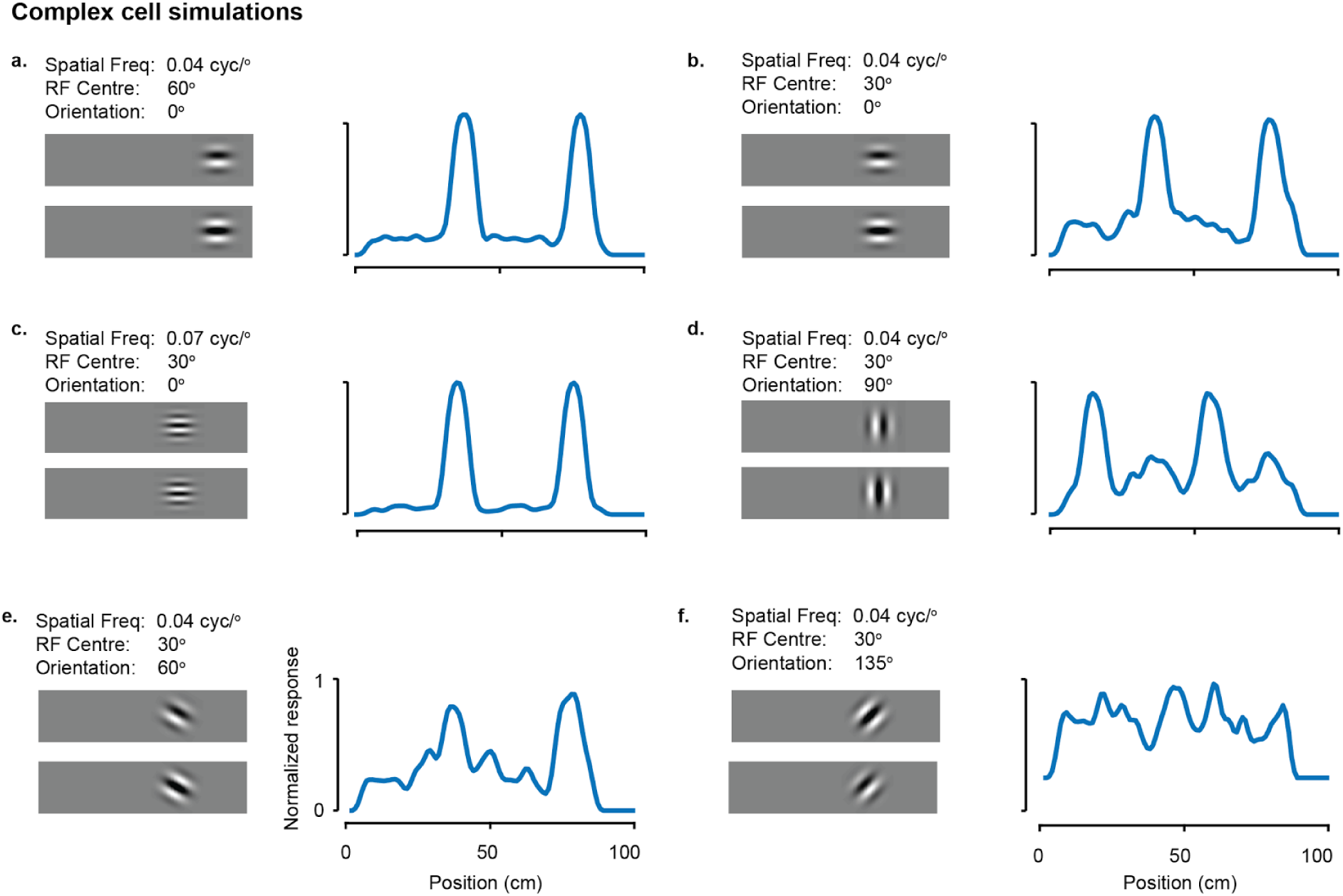
Simulation of purely visual responses to position in VR. Responses of 6 simulated neurons, with purely visual responses produced by a complex cell model, with varying spatial frequency, orientation, or receptive field centres. The images on the left of the panel show the quadrature pair of complex cell filters, and on the right is the response of the simulation of that pair as a function of position in the virtual environment. Simulated complex cells had spatial frequencies, and orientations that are commonly observed in mouse V1 (sf: 0.04, 0.05, 0.06 or 0.07 cyc/°; orientations: 0^°^ to 90^°^ with twice more cells for cardinal orientations). The receptive field had positions > 40^°^ azimuth (40^°^, 50^°^, 70^°^, 80^°^), similar to the V1 neurons we considered for analysis. In rare cases (like in **f**) when the receptive fields do not match the features of the environment, we get little selectivity along the corridor. These cases lead to lower value for the response ratios.

**Supplementary Figure 4:**
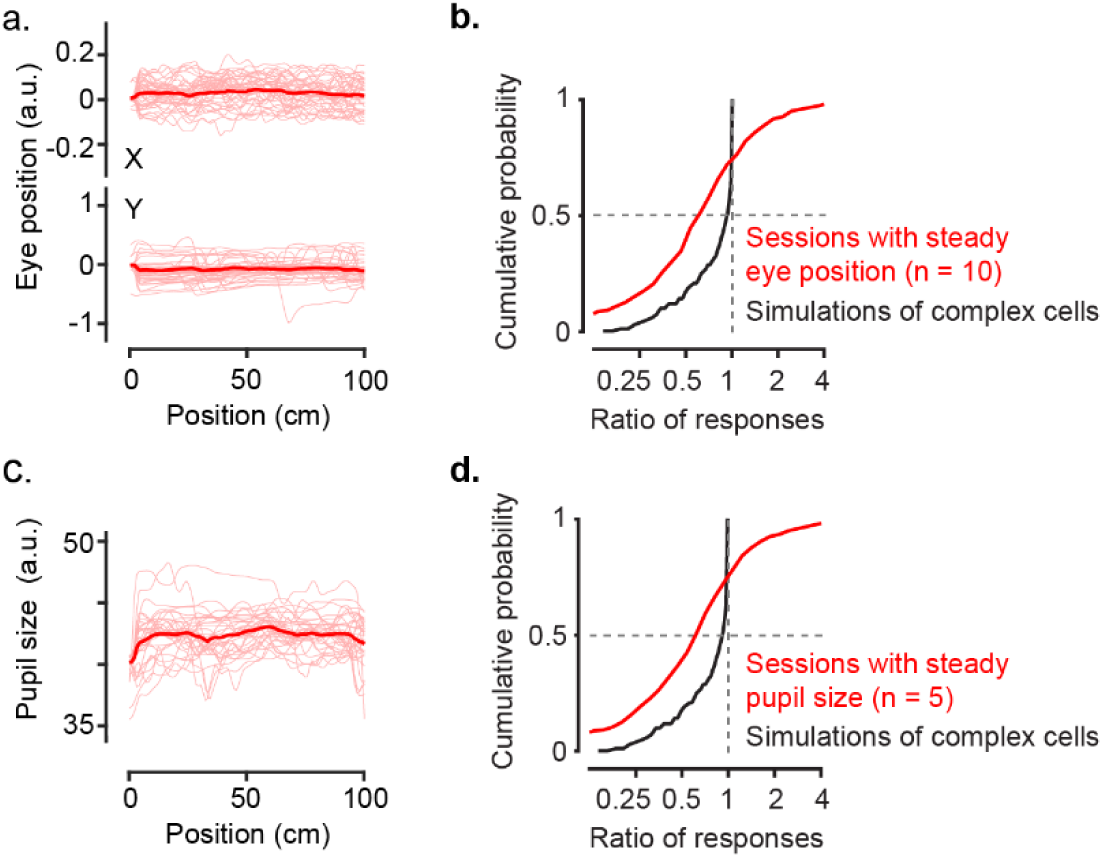
The spatial modulation of V1 responses is not a result of pupil position or diameter. **a,** Pupil position as a function of location in the virtual corridor, for an example session where the position of the pupil was on average the same along the corridor. Thin red curves: position trajectories on individual trials; thick curves, average. Top and bottom panels: x- and y-coordinates of the pupil. **b,** Distribution of response ratios for sessions with steady eye position (10 sessions; *red*) and for simulations of complex cells *(black)*.The two distributions are significantly different (two-sample Kolmogorov-Smirnov test; p≪0.001). **c,** Pupil size as a function of position for an example session where pupil size was on average the same along the VR corridor. **d,** Distribution of response ratios for sessions with steady pupil size (5 sessions; *red)* and for simulations of complex cells *(black).* The two distributions are significantly different (two-sample Kolmogorov-Smirnov test; p≪0.001).

**Supplementary Figure 5:**
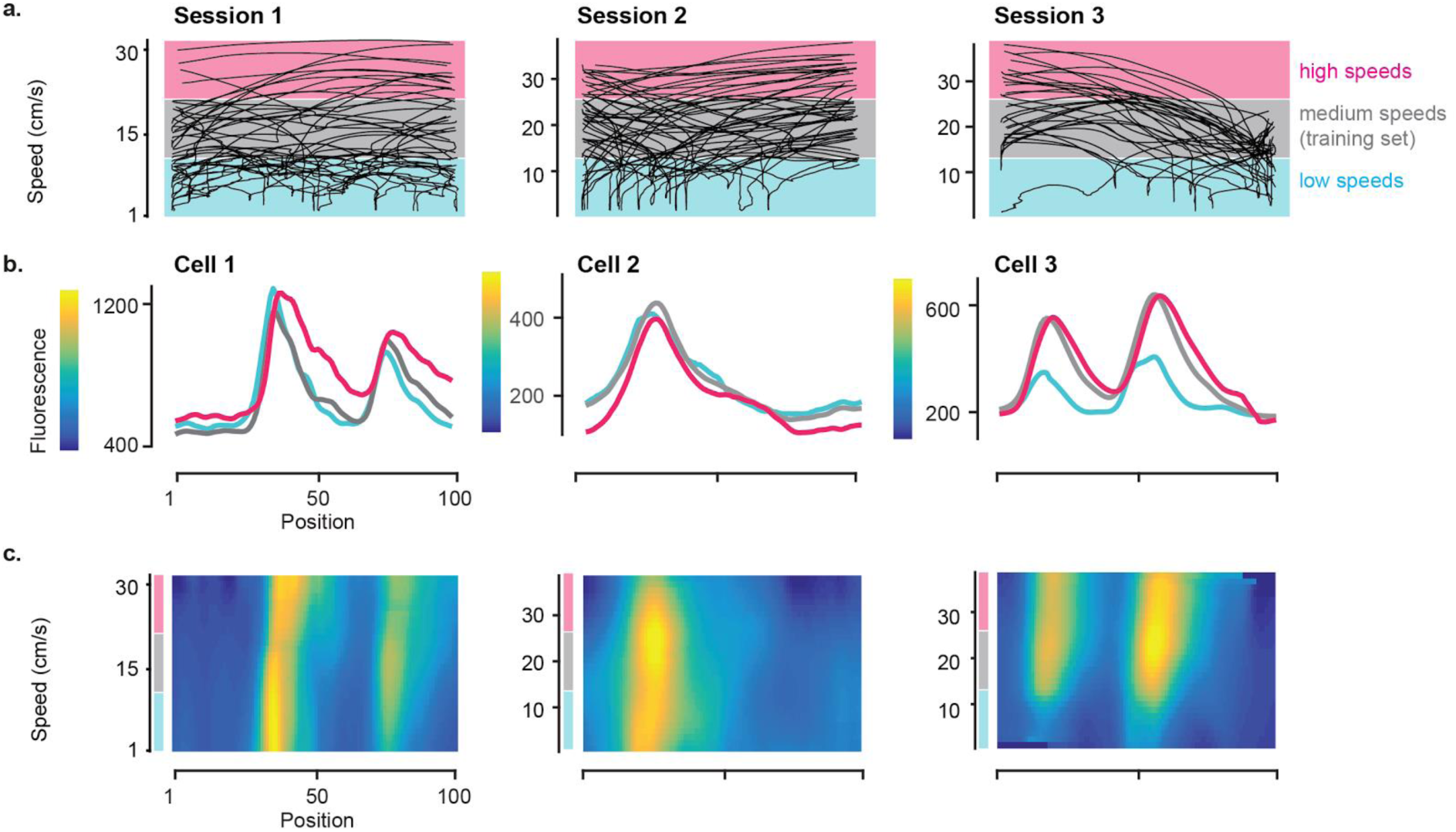
The spatial modulation of responses in V1 cannot be explained by speed. **a,** Single-trial trajectories of speed as a function of position in the virtual reality environment for three example recording sessions. **b,** Response profile of an example V1 cells in each session as a function of position in the room for three different speed ranges, corresponding to the 3 shading bands in **a**. **c,** Two-dimensional response profiles of the same example neurons showing activity as a function of position and running speed for speeds higher than 1cm/s.

**Supplementary Figure 6:**
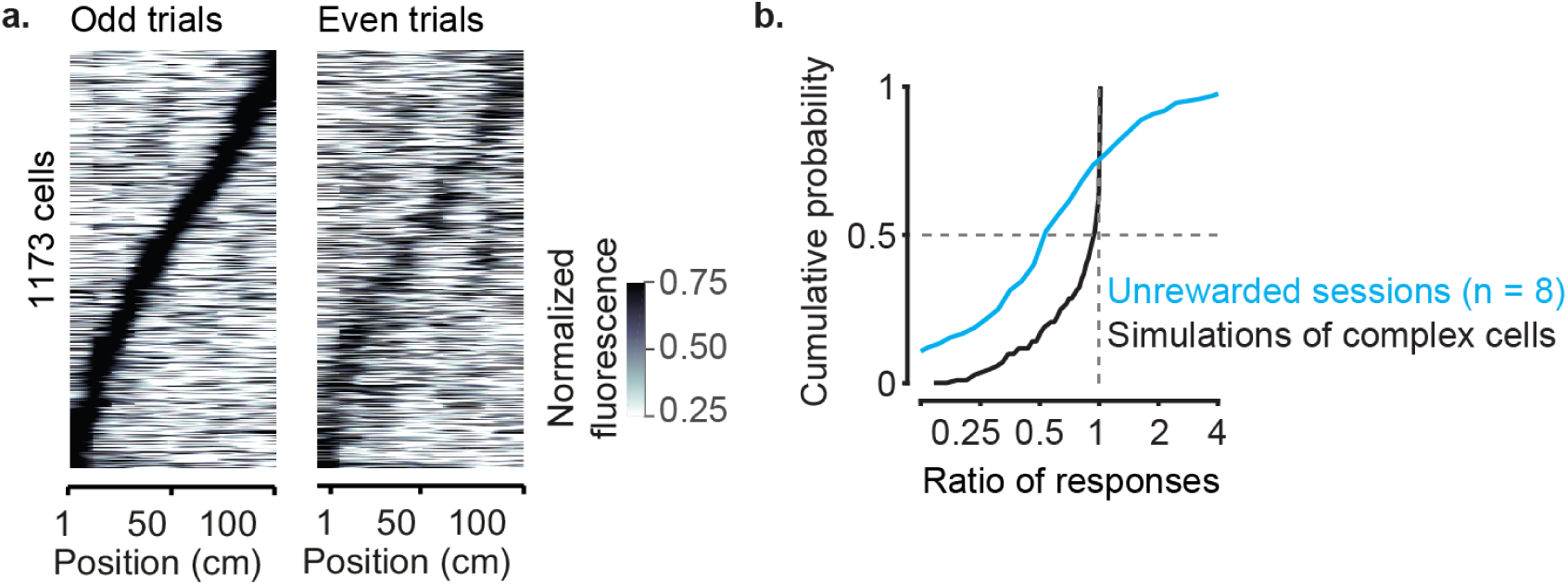
The spatial modulation of V1 responses cannot be explained by reward. **a,** Normalized response as a function of position in the virtual corridor, across sessions without reward (1173 cells). Data come from 2 out 4 mice that freely ran the environment without reward (8 sessions). Responses in even trials *(right)* are ordered according to the position of maximum activity measured in odd trials *(left)*. **b,** Distribution of response ratios for unrewarded sessions (8 sessions; *cyan*) and for simulations of complex cells *(black)*. The two distributions are significantly different (two-sample Kolmogorov-Smirnov test; p≪0.001).

**Supplementary Figure 7:**
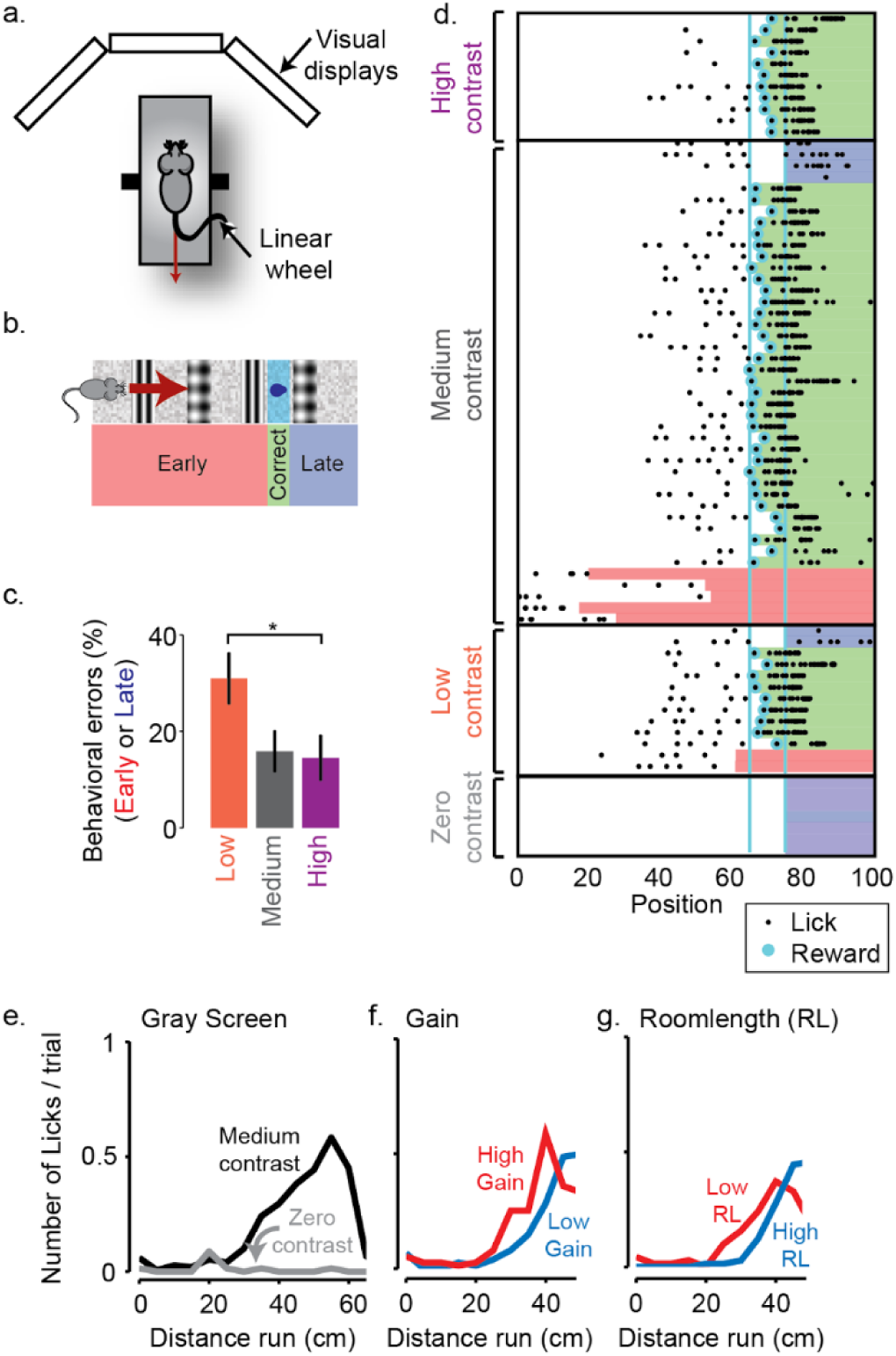
Behavioural performance in the task. **a,** Cartoon of the experimental setup. Virtual reality was presented on three visual displays. **b,** Illustration of the virtual reality environment with four prominent landmarks, a reward zone, and the zones that define trial types: *Early, Correct* and *Late.* **c,** Percentage of trials during which the animal makes behavioural errors, by licking either too early or too late at three different contrast levels: 18% (low), 60% (medium) or 72% (high). **d,** Illustration of performance on all trials of one example recording session. Each row represents a trial, *black dots* represent positions where the animal licked, and *cyan dots* indicate the delivery of a water reward. *Coloured bars* indicate the outcome of the trial, *red: Early, green: Correct, blue: Late.* **e-g,** Successful performance relies on vision: **e,** The animal does not lick when the room was presented at zero contrast. **f**. If the gain between the animals’ physical movement and movement in the virtual environment was increased, the animals licked after running a shorter physical distance. **g,** If the position of the visual cues was shifted forward or back (high/low room length (RL)), the lick position shifted accordingly, indicating that the animals rely on vision to perform the task.

## Methods

All experiments were conducted according to the UK Animals (Scientific Procedures) Act, 1986 under personal and project licenses issued by the Home Office following ethical review.

For simultaneous recordings in V1 and CA1, we used four C57BL/6 mice (all male, implanted at 4-8 weeks of age). For calcium imaging experiments, we used double or triple transgenic mice expressing GCaMP6 in excitatory neurons (3 females, 1 male, implanted at 4-6 weeks). The double transgenic expressed GCaMP6 slow (Wekselblatt et al., 2016) (Camk2a-tTA;tetO-G6s). The triple transgenics expressed GCaMP6 fast (Madisen et al., 2015) (Emx1-Cre;Camk2a-tTA;Ai93, 3 mice). None of these mice displayed the aberrant activity that is sometimes seen in Ai93 mice (Steinmetz et al., 2017).

### Virtual Environment and task

The virtual reality environment was a corridor adorned with a white noise background and four landmarks: two grating stimuli oriented orthogonal to the corridor and two plaid stimuli (Figure 1**a**). The corridor dimensions were 100 × 8 × 8 cm, and the landmarks (8 cm wide) were centred 20, 40, 60 and 80 cm from the start of the corridor. The animal navigated the environment by walking on a custom-made polystyrene wheel (15 cm wide, 18 cm diameter). Movements of the wheel were captured by a rotary encoder (2400 pulses/rotation, Kübler, Germany), and used to control the virtual reality environment presented on three monitors surrounding the animal, as previously described(Saleem et al., 2013). When the animal reached the end of the corridor, it was placed back at the start of the corridor after a 3-5 s presentation of a grey screen. Trials longer than 120 s were timed out and were excluded from further analysis.

Mice used for calcium imaging ran freely through the corridor, with no specific task. Two of the mice were motivated to run with water rewards. One animal received rewards at random positions along the corridor. The other received rewards at the end of the corridor. To control for the effect of the reward on V1 responses, no reward was delivered in some sessions (n = 8 sessions; 2 animals).

Mice used for simultaneous V1-CA1 recordings were trained to lick in a specific region of the corridor, the reward zone. This zone was centred at 70 cm and was 8 cm wide. Trials in which the animals were not engaged in the task, i.e. when they ran through the environment without licking, were excluded from further analysis. The animal was rewarded for correct licks with ~2 μl water using a solenoid valve (161T010; Neptune Research, USA), and licks were monitored using a custom device that detected breaks in an infrared beam.

### Surgery and training

The surgical methods are similar to those described previously (Ayaz et al., 2013; Saleem et al., 2013). Briefly, a custom head-plate with a circular chamber (3-4 mm diameter for electrophysiology; 8 mm for imaging) was implanted on 4-8 week mice under isoflurane anaesthesia. For imaging, we performed a 4 mm craniotomy over left visual cortex by repeatedly rotating a biopsy punch. The craniotomy was shielded with a double coverslip (4 mm inner diameter; 5 mm outer diameter). After 4 days of recovery, some mice were water restricted (> 40 ml / kg / day) and were trained for 30-60 min, 5-7 days/week.

Mice used for simultaneous V1-CA1 recordings were trained to lick selectively in the reward zone using a progressive training procedure. Initially, the animals were rewarded for running past the reward location on all trials. After this, we introduced trials where the animal was rewarded only when it licked in the rewarded region of the corridor. The width of the reward region was progressively narrowed from 30 cm to 8 cm across successive days of training. To prevent the animals from licking all across the corridor, trials were terminated early if the animal licked more than a certain number of times before the rewarded region. We reduced this number as the animals performed more accurately, typically reaching a level of 4-6 licks by the time recordings were made. Once a sufficient level of performance was reached, we controlled on some (random) trials that the animal performed the task visually by measuring the performance when decreasing the visual contrast or changing the distance to the reward zone (Supplementary Figure 7). Training was carried out for 3-5 weeks. Animals were light shifted (9 am light off, 9 pm light on) and experiments were performed during the day.

### Widefield calcium imaging

For widefield imaging we used a standard epi-illumination imaging system (Ratzlaff and Grinvald, 1991; Carandini et al., 2015) together with an SCMOS camera (pco.edge, PCO AG). A Leica 1.6x Plan APO objective was placed above the imaging window and a custom black cone surrounding the objective was fixed on top of the headplate to prevent contamination from the monitors’ light. The excitation light beam emitted by a high-power LED (465 nm LEX2-B, Brain Vision) was directed onto the imaging window by a dichroic mirror designed to reflect blue light. Emitted fluorescence passed through the same dichroic mirror and was then selectively transmitted by an emission filter (FF01-543/50-25, Semrock) before being focused by another objective (Leica 1.0 Plan APO objective) and finally detected by the camera. Images of 200 x 180 pixels, corresponding to an area of 6.0 × 5.4 mm were acquired at 50 Hz.

To measure retinotopy we presented a 14^°^-wide vertical window containing a vertical grating (spatial frequency 0.15 cycles/deg), and swept (Kalatsky and Stryker, 2003; Yang et al., 2007) the horizontal position of the window over 135^°^ of azimuth angle, at a frequency of 2 Hz. Stimuli lasted 4 s and were repeated 20 times (10 in each direction). We obtained maps for preferred azimuth by combining responses to the 2 stimuli moving in opposite direction, as previously described (Kalatsky and Stryker, 2003).

### Two-photon imaging

Two-photon imaging was performed with a standard multiphoton imaging system (Bergamo II; Thorlabs) controlled by ScanImage4 (Pologruto et al., 2003). A 970 nm laser beam, emitted by a Ti:Sapphire Laser (Chameleon Vision, Coherent), was targeted onto L2/3 neurons through a 16x water-immersion objective (0.8 NA, Nikon). Fluorescence signal was transmitted by a dichroic beamsplitter and amplified by photomultiplier tubes (GaAsP, Hamamatsu). The emission light path between the focal plane and the objective was shielded with a custom-made plastic cone, to prevent contamination from the monitors’ light. In each experiment, we imaged 4 planes set apart by 40 μm. Multiple-plane imaging was enabled by a piezo focusing device (P-725.4CA PIFOC, Physik Instrumente), and an electro-optical modulator (M350-80LA, Conoptics Inc.) which allowed adjustment of the laser power with depth. Images of 512x512 pixels, corresponding to a field of view of 500×500 μm, were acquired at a frame rate of 30 Hz (7.5 Hz per plane).

Pre-processing of raw imaging movies involved 1) image registration to correct for brain movement, 2) ROI extraction, i.e. cell detection and 3) correction for neuropil contamination. Image registration, cell detection and neuropil signal extraction were performed with the Suite2p pipeline (Pachitariu et al., 2016). Cells with noisy baseline or extremely seldom firing were excluded from further analysis.

For neuropil correction, we used an established method (Peron et al., 2015; Dipoppa et al., 2016). We used Suite2p to determine a mask surrounding each cell’s soma, the ‘neuropil mask’. The inner diameter of the mask was 3 μm and the outer diameter was < 45 μm. For each cell we obtained a correction factor, α, by regressing the binned neuropil signal (20 bins in total) from the 5^th^ percentile of the raw binned cell signal. For a given session, we obtained the average correction factor across cells. This average factor was used to obtain the corrected individual cell traces, from the raw cell traces and the neuropil signal, assuming a linear relationship. All correction factors fell within 0.7 and 0.9.

### Pupil tracking

We tracked the eye of the animal using an infrared camera (DMK 21BU04.H, Imaging Source) and a zoom lens (MVL7000, Navitar) at 25 Hz. Pupil position and size were calculated by fitting an ellipsoid to the pupil for each frame using a custom software. X and Y positions of the pupil were derived from the centre of mass of the fitted ellipsoid.

### Electrophysiological recordings

On the day prior to the first recording session, we made two 1 mm craniotomies, one over CA1 (1.0 mm lateral, 2.0 mm anterior from lambda), and a second one over V1 (2.5 mm lateral, 0.5 mm anterior from lambda). We covered the chamber using KwikCast (World Precision Instruments) and the animals were allowed to recover overnight. The CA1 probe was lowered until all shanks were in the pyramidal layer. We waited ~30 min for the tissue to settle before starting the recordings. In two animals, we dipped the probes in red-fluorescent DiI (Figure 2**a**). In these animals, we had only one recording session. The other animals had two and four recording sessions.

Offline spike sorting was carried out using the KlustaSuite (Rossant et al., 2016) package, with automated spike sorting using KlustaKwik (Kadir et al., 2014), followed by manual refinement using KlustaViewa (Rossant et al., 2016). Hippocampal interneurons were identified based on their spike time autocorrelation and excluded from further analysis. Only time points with running speeds greater than 5 cm/s were included in further analyses.

### Analysis of response profiles for electrophysiological data

To calculate the response profile of each neuron as a function of position in the virtual corridor we used a local smoothing method (Loader, 1996; Harris et al., 2002, 2003). We first smoothed the firing rate using a 250 ms Gaussian window. We then discretized the position of the animal in 2 cm bins, yielding 50 bins and we calculated the spike count map and occupancy map for each neuron. Both the spike count and occupancy maps were smoothed by convolving them with a common Gaussian window whose width was optimized to maximize reliability (see below), and the response profile was calculated as the ratio of the smoothed spike count map and the occupancy map (Saleem et al., 2013).

Response profile reliability was calculated as the fraction of variance in firing rate explained by the response profile:

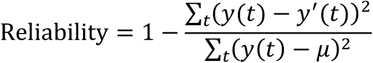

where *y(t)* is the firing rate of the neuron at time *t, y’(t)* was the prediction by the place-field for the same time bin and *μ* is the mean firing rate of the training data(Saleem et al., 2013). We used five-fold cross-validation to calculate place fields and reliability. Only neurons with a reliability greater than 0.01 were considered for further analysis.

### Analysis of response profiles for two-photon data

To obtain response profiles as a function of position along the corridor, we averaged the neuropil-corrected activity of each cell in 1-cm-wide bins (100 bins in total) and smoothed with a 5cm Gaussian window. Only time points with running speeds greater than 1 cm/s were included in further analyses. For consistency with the response profiles obtained from electrophysiological data, we only looked at responses for which the cross-validated reliability was higher than 0.01. These cells were considered to have activity significantly modulated by position in the corridor. To model single-cell activity under the assumption that responses are identical in the two segments of the corridor, we fit (using least squares) a model function to the response profile along the visually-matching segment where the cell peaked. The model function was the sum of two Gaussians that meet at the peak. To obtain a prediction along the whole corridor, we then duplicated the fitted response at ± 40 cm away from the maximum. Cells which had a maximal response too close to the start or the end of the corridor (0-15 cm or 85-100cm) were not considered for analysis of the ratio of responses. This excluded cells which responded too close to the start or the end of the corridor, which were outside the visually-matching segments Two-dimensional response profiles with respect to position and speed (Supplementary Figure 5c) were calculated as previously described (Saleem et al., 2013).

### Decoding population activity & using position decoded from other cells

Population activity was decoded using an independent Bayes decoder. For every time bin, we calculated the probability of being at a location *x* given population response *R* as:

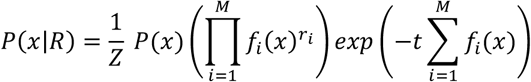

where *f_i_(x)* is the response profile and *r_i_* is the spike count of the *i^th^* neuron in a time bin, *M* is the number of neurons and *t* is the time window. *Z* is a normalizing constant, which makes the probabilities across all positions sum to one (Zhang et al., 1998; Bendor and Wilson, 2012). The probability of being in the reward zone was calculated by summing the posterior probabilities in the reward zone, and normalized relative to the value in correct trials (Figure 3c and 3f). When calculating joint distributions (Figures 2 and 3), we smoothed the distribution by a Gaussian window with a width of 4 spatial bins. To account for the effects of position and speed on calculating the correlations between V1 and CA1 decoding errors, we shuffled the data within the time points when the animal was at the same position (within 2cm) and ran in a specific speed range (5 cm/s bins: 5-10 cm/s to 30-35 cm/s).

### Simulation of V1 complex cells

Response profiles expected from purely visual neurons were obtained from simulations of a population of complex receptive fields. Complex receptive fields were modelled as two Gabor filters in spatial quadrature (i.e. shifted in spatial phase by 90 deg) having the same orientation and spatial frequency. Responses were simulated by convolving the VR images at successive positions along the corridor with the pair of Gabor filters and taking the sum of their squared outputs (energy model (Movshon et al., 1978; Carandini, 2006)). The receptive fields were designed so to simulate different orientation selectivity (from 0^°^, 15^°^,…, 165^°^; we overrepresented the cardinal orientations) spatial frequency selectivity (0.04, 0.05, 0.06 and 0.07 cycles/deg), which are the ranges typically observed in the mouse visual system (Niell and Stryker, 2010; Andermann et al., 2011; Marshel et al., 2011). The receptive fields were simulated to cover azimuths from 40^°^ to 80^°^, matching the receptive field position of the cells we focused on in our recordings.

